# Branch-resolved IFN/STING-linked lesion-state architecture in psoriasis: multi-cohort derivation, held-out bulk replication and single-cell context analysis

**DOI:** 10.64898/2026.07.19.739418

**Authors:** Hsiao-Yu Sun, Tien-Lin Chang, Zhi-Hong Wen, Hsi-Wen Sun

## Abstract

**Objective and design:** We tested whether a frozen branch-resolved IFN/STING-linked lesion-state architecture derived from paired psoriasis transcriptomes would transport to an untouched bulk RNA-sequencing cohort and remain detectable within broad cell compartments.

**Material or subjects:** Seven paired bulk cohorts formed the derivation backbone. GSE121212 provided held-out replication using 27 matched lesional/nonlesional pairs; treatment-facing cohorts and GSE228421 provided pharmacodynamic and cellular-context analyses.

**Treatment:** No intervention was administered by the authors; public datasets included ustekinumab-, etanercept-, secukinumab- and risankizumab-exposed samples.

**Methods:** Frozen modules and branches were scored after cohort-wide gene standardization. GSE121212 counts underwent trimmed mean of M-values normalization and log2 counts-per-million transformation. Paired effects, upper-quartile STING-high contrasts, lesional Spearman coupling and prespecified rank concordance were evaluated.

**Results:** GSE121212 lesional-minus-nonlesional effects were 1.288 for STING-core, 1.478 for IFN-responsive activity and 1.188 for proximal-only STING; all lower 95% confidence limits exceeded zero. Coupling-vector concordance was strong (rho = 0.893, P = 0.0068), whereas STING-high effect concordance was moderate (rho = 0.536), yielding partial replication.

**Conclusions:** Held-out replication was partial: the IFN/STING lesion anchor and proximal/IFN-dominant coupling transported, while broader ordering did not fully transport. Transcriptomic findings do not establish biochemical STING activation, causality or clinical thresholds.

## Introduction

Psoriasis is a chronic immune-mediated skin disease in which biologics targeting tumour necrosis factor (TNF), interleukin (IL)-23 and IL-17 have transformed clinical control. However, partial response, relapse and incomplete molecular normalization remain important clinical and translational problems [1–4]. These limitations do not weaken the central role of the IL-23/Th17 axis; instead, they suggest that plaque-level inflammatory states may not be fully captured by a single downstream cytokine measurement.

Human transcriptomic studies have repeatedly identified immune, keratinocyte, stromal and vascular abnormalities in psoriatic lesions [5–8]. Existing pathway analyses often report interferon, NF-κB, IL-36, IL-17, keratinocyte differentiation and myeloid/neutrophil programmes as parallel outputs. More recent molecular-stratification and treatment-response studies further support lesion-level heterogeneity [9–13]. What remains less clear is whether these outputs can be organized into a reproducible lesion-state architecture that distinguishes target-proximal, response-facing, effector and context branches.

The cyclic GMP-AMP synthase-stimulator of interferon genes (cGAS-STING) pathway links cytosolic DNA sensing to TBK1/IRF3-driven type I interferon and NF-κB-associated responses [14, 15]. STING-linked and type I interferon biology has been implicated in psoriasiform inflammation, dendritic-cell amplification, wound-associated initiation, IL-36/IL-23 crosstalk and neutrophil/NETosis-related pathology [16–23]. Nevertheless, transcriptomic evidence should not be interpreted as biochemical STING activation unless supported by orthogonal protein or functional assays.

We therefore restructured the analysis around a conservative premise: known psoriasis inflammatory biology can be converted into a frozen, branch-resolved lesion-state framework and tested for derivation-set reproducibility, transport to an untouched bulk cohort, treatment-linked remodeling and broad-cell-type context. Unlike conventional pathway-enrichment analyses that examine signatures separately, the frozen framework required the relationships among branches to survive an external transport test. The objective was not to claim discovery of a previously unknown psoriasis pathway or to establish STING as a causal driver. Rather, we asked which parts of an IFN/STING-linked lesion-state architecture remain reproducible across distinct datasets and analytical scales (Fig. 1).

**Figure 1.**
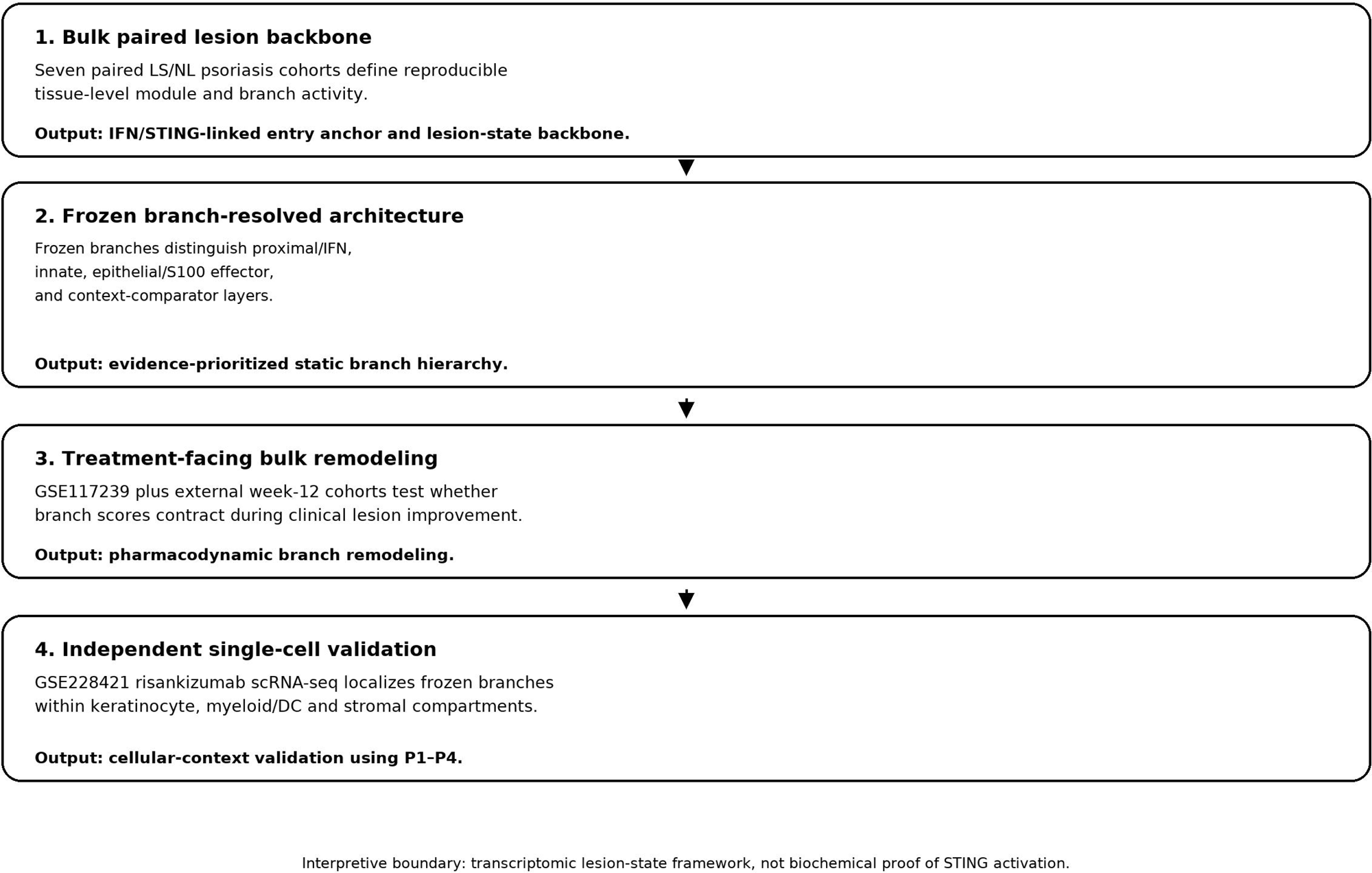
Integrated study design. Seven paired bulk cohorts define the derivation architecture; treatment-facing cohorts test pharmacodynamic remodeling; untouched GSE121212 tests held-out bulk transport; and GSE228421 provides broad-cell-type context and major-composition robustness. The study separates derivation, held-out replication, treatment support and cellular-context roles.

## Materials and methods

### Study design and public datasets

This study was a secondary analysis of public human psoriasis transcriptomic datasets from the Gene Expression Omnibus (GEO) [24, 25]. Seven paired lesional/nonlesional bulk cohorts formed the derivation backbone: GSE6710, GSE13355, GSE14905, GSE30999, GSE34248, GSE41662 and GSE53552 [26–32]. GSE78097 provided severity context [33]. GSE117239 provided the primary treatment-response layer, and GSE11903 and GSE171012 provided external week-12 therapy-facing analyses [34–36]. GSE121212 was reserved as an untouched held-out bulk RNA-sequencing replication cohort [37]. GSE228421 provided longitudinal single-cell context analysis during risankizumab treatment [42, 43]. Cohort roles, retained counts and analysis roles are summarized in Online Resource 1.

Analyses were designed around frozen module and branch definitions rather than data-adaptive gene selection in the validation datasets. The seven paired bulk cohorts defined the tissue-level lesion architecture, cross-cohort reproducibility and frozen discovery vectors. GSE121212 tested transport of the prespecified lesion anchor and branch architecture without revising gene sets or success criteria after outcome inspection. GSE228421 tested broad-cell-type-stratified distribution and robustness against an explanation based solely on shifts among major annotated cell types; it was not treated as an independent replication of the complete bulk hierarchy.

### Branch definitions and bulk scoring

Primary modules included a STING-core module, a type I interferon module and an IL-23/Th17 proxy. Downstream branches included proximal bridge, IFN-responsive, NF-κB broad innate, keratinocyte effector, myeloid/neutrophil proxy, immune proxy and keratinocyte differentiation branches. The IFN-responsive branch used the same gene set as the type I interferon entry module. The distinction was functional rather than genetic: the entry module tested lesion-level enrichment across cohorts, whereas the branch was used within the frozen branch framework to test STING-aligned preferential activation, treatment-linked reversibility and cellular localization. Each sample was scored using within-cohort z-mean scoring across genes available on the corresponding platform (Online Resource 2).

Paired lesional-minus-nonlesional effects were estimated within cohorts and summarized using random-effects meta-analysis with restricted maximum likelihood [41]. Within-lesion coupling was estimated between STING-core activity and other modules or branches. STING-high analyses compared top-quartile STING-core lesions with comparator lesions within each cohort. Proximal-only STING sensitivity analysis removed downstream interferon-output genes from the STING reference to reduce circularity.

### Held-out GSE121212 bulk RNA-sequencing replication

GSE121212 contains 147 skin transcriptomes, including 55 psoriasis samples from 28 patients [37]. Before scoring, the official count matrix and GEO metadata were checksum-frozen, sample identities were resolved deterministically, and 27 patients with complete matched lesional/nonlesional samples were retained for paired inference. The unmatched lesional sample from PSO_034 was excluded from paired analyses but retained among 28 lesions for prespecified lesion-only STING-high and coupling analyses.

The primary replication lane summed duplicate gene-symbol rows, removed only genes with zero counts in all 147 samples, applied edgeR trimmed mean of M-values normalization and log2 counts-per-million transformation with prior count 2 [38], and standardized each gene across all 147 samples. Frozen module and branch scores were means of available gene z scores using the original minimum-retention rules. A legacy sensitivity lane averaged duplicate-symbol rows and applied raw-count gene-wise standardization to mirror the earlier GSE171012 implementation.

The frozen replication hierarchy comprised three components. Anchor replication required positive paired STING-core and IFN-responsive effects with lower 95% confidence limits above zero, together with a positive proximal-only STING effect. Architecture transport compared the seven-branch STING-high effect and lesional coupling vectors with the frozen discovery vectors using Spearman rank correlation and prespecified direction/ranking criteria. Overlap-robust analysis used proximal-only STING as the reference. Patient-level paired effects used t-based 95% confidence intervals; STING-high contrasts used the upper quartile among 28 lesions; coupling used two-sided Spearman correlation. The primary TMM-log2CPM lane was publication-authoritative, and the raw-count lane was sensitivity analysis only (Online Resource 6).

### Treatment-facing bulk analyses

In GSE117239, baseline branch scores and week-12-minus-baseline changes were summarized by PASI75 response status. Branch-level analyses included responders and nonresponders with paired week-12 data. External GSE11903 and GSE171012 analyses tested patient-level baseline-to-week-12 changes against zero. These analyses were interpreted as treatment-linked pharmacodynamic remodeling and not as proof of causal pathway targeting. Detailed bulk branch architecture, treatment and robustness statistics are provided in Online Resource 3.

### Composition-aware and gene-level robustness analyses

To reduce the risk that branch associations reflected only tissue composition, composition-aware models adjusted STING-core-to-branch relationships using immune-proxy and keratinocyte-differentiation context scores. Representative-gene analyses assessed consistency of selected genes across paired lesion comparisons, STING-high analyses and lesion-only associations. These analyses were intended as robustness and anchoring checks, not as independent mechanistic validation.

### GSE228421 single-cell context analysis

GSE228421 contains longitudinal single-cell RNA sequencing of skin biopsies from individuals with severe psoriasis receiving risankizumab [42, 43]. The downloaded raw 10x archive was audited before analysis. Seventeen samples with complete barcodes, features and matrix files were retained. Four patients had complete paired baseline lesional/nonlesional samples and complete lesional day-0, day-3 and day-14 trajectories; one additional day-0 lesional sample was retained only for descriptive visualization. Single-cell filtering, annotation and cell-type-count details are provided in Online Resource 4.

Because the downloaded matrices represented raw droplet-level barcode matrices rather than filtered cell matrices, each sample was filtered independently before concatenation. The primary filter retained barcodes with n_genes_by_counts at least 300, total_counts at least 1000 and mitochondrial percentage at most 20%. The filtered object contained 166,954 cells and 36,601 genes. Counts were normalized to 10,000 per cell and log1p transformed before branch scoring.

Frozen branch gene sets were transferred without re-selection. For each cell, each branch score was computed as the mean log1p-normalized expression of available branch genes. Branch gene coverage was complete for most signatures; STING-core retained 9 of 10 genes and proximal-only STING retained 5 of 6 genes because MB21D1 was absent from the processed feature set.

### Clustering, annotation and cell-type summaries

Highly variable genes were used for principal component analysis, nearest-neighbor graph construction, Leiden clustering and uniform manifold approximation and projection visualization using Scanpy [44]. Broad cell-type labels were assigned manually using individual canonical markers for keratinocytes, fibroblasts, endothelial cells, pericyte/smooth-muscle cells, melanocytes, T/NK cells, myeloid/DC and Langerhans/DC cells, mast cells and B/plasma cells. Langerhans/DC clusters were retained within the broad myeloid/DC compartment for primary summaries rather than analysed as a separate primary stratum. Low-confidence mixed clusters were excluded from primary cell-type-stratified inference (Online Resource 4).

Branch scores were summarized by patient, sample and broad cell type. Patients, not cells, were the unit of inference. Baseline analyses compared paired lesional and nonlesional day-0 scores within P1-P4. Longitudinal analyses compared lesional day-3 and day-14 scores with day-0 within P1-P4. P values were interpreted descriptively because of the small patient-level sample size and the cellular-context role of this analysis. Persistence of directional shifts within broad annotated cell types was interpreted as arguing against an explanation based solely on shifts among major cell-type proportions; it does not exclude finer within-compartment composition or cell-state changes. Full matrices are provided in Online Resource 5.

### Reproducibility and software

Bulk analyses were scripted in R using GEOquery, limma, edgeR, GSVA and metafor where applicable [25, 38–41]. Single-cell analyses were scripted in Python using Scanpy, AnnData, NumPy, pandas, SciPy and statsmodels [44]. Summary tables, figure-source matrices, frozen-design documents, checksums and workflow logs were retained for auditability. Curated scripts, run-order notes and reporting notes are described in Online Resource 7. Artificial intelligence use was limited to editorial language support; see Declarations and Online Resource 7.

## Results

### A reproducible IFN/STING entry anchor is present in psoriasis lesions

Across seven paired bulk cohorts, lesional skin showed positive enrichment of all three prespecified entry modules (Fig. 2; Online Resource 3). The pooled lesional-minus-nonlesional effect was largest for the type I interferon module (delta = 1.347, 95% CI 1.207 to 1.487; P = 4.83 x 10^-79), followed by STING-core (delta = 0.807, 95% CI 0.679 to 0.935; P = 2.72 x 10^-35) and the IL-23/Th17 proxy (delta = 0.639, 95% CI 0.520 to 0.757; P = 4.15 x 10^-26).

**Figure 2.**
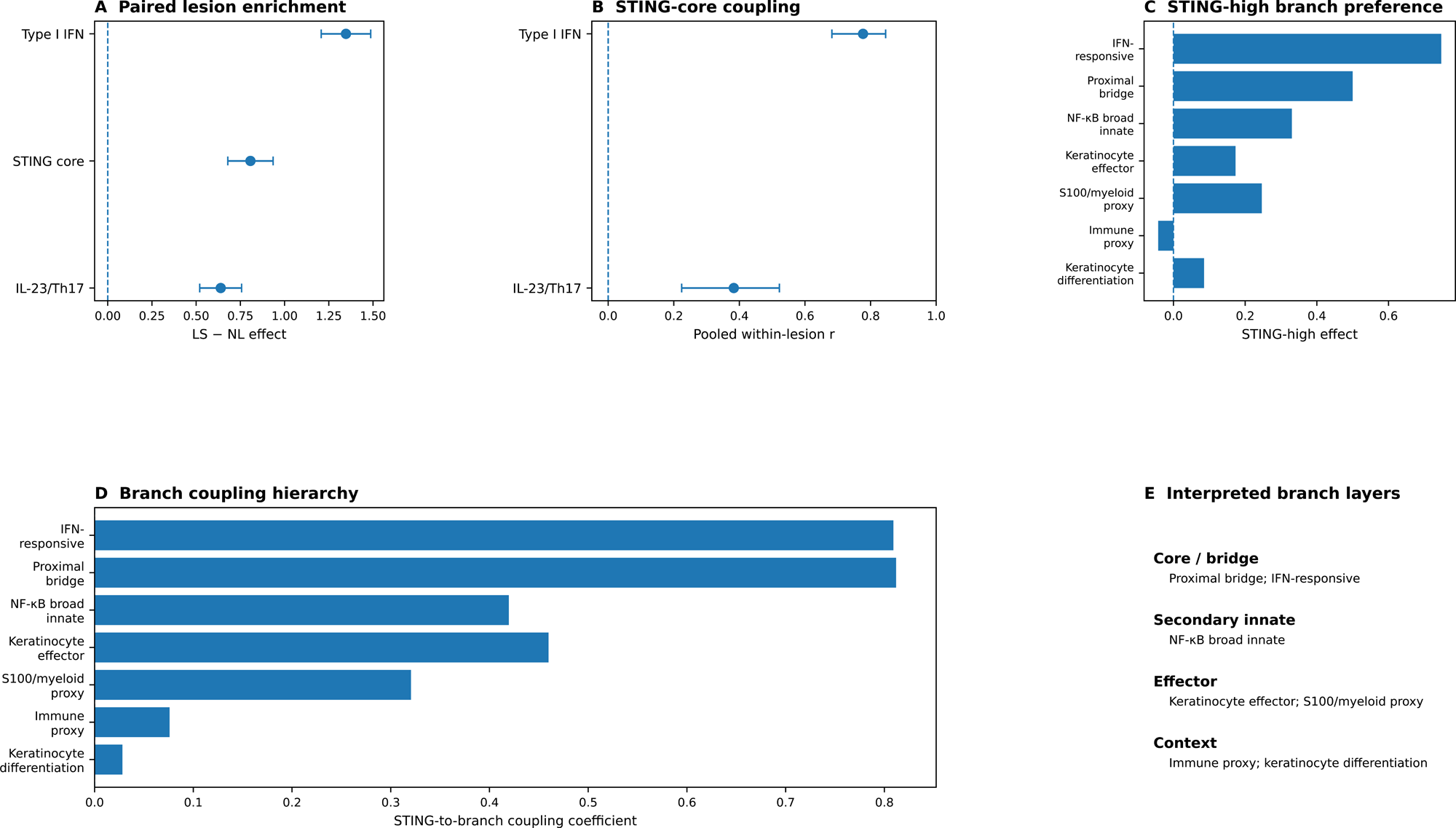
Bulk IFN/STING entry anchor and branch hierarchy. Paired lesional-minus-nonlesional effects, within-lesion STING-core coupling and STING-high stratification identify a proximal/IFN core, secondary innate layer and downstream effector branches. Branch ordering reflects reproducibility, STING-high preference, coupling and treatment-linked reversibility; it does not establish causal STING activation.

Within lesions, STING-core coupled more strongly with type I interferon activity than with the IL-23/Th17 proxy. The pooled correlation was 0.777 (95% CI 0.682 to 0.846) for type I interferon and 0.383 (95% CI 0.224 to 0.522) for the IL-23/Th17 proxy. STING-high lesions also had higher interferon and IL-23/Th17 proxy activity than comparator lesions, establishing IFN/STING activity as an entry anchor rather than the final endpoint.

### Severity-context analysis does not show a monotonic IFN/STING gradient

In GSE78097, severe compared with mild lesions did not show higher IFN/STING activity. The severe-minus-mild differences were -0.238 for STING-core (95% CI -0.489 to 0.013) and -0.590 for IFN-responsive activity (95% CI -1.158 to -0.022), while the keratinocyte proxy was higher (0.615, 95% CI 0.192 to 1.038) and the myeloid/neutrophil proxy was lower (-0.303, 95% CI -0.583 to -0.023). These findings indicate that the lesion-associated IFN/STING anchor does not scale monotonically with cross-sectional clinical severity and that severity-associated tissue context differs from the lesional-versus-nonlesional contrast (Online Resource 3).

### STING-high lesions resolve into a branch hierarchy rather than uniform inflammation

STING-high lesions did not show nonselective elevation of all inflammatory readouts. IFN-responsive and proximal-bridge branches showed the strongest STING-high preference and STING-to-branch coupling, with STING-high effects of 0.747 and 0.500 and pooled STING-core coupling coefficients of 0.809 and 0.812, respectively (Fig. 2; Table 1; Online Resource 3).

**Table 1.** Integrated branch-resolved IFN/STING-linked architecture, held-out replication and proposed analytical roles.

| Branch | Proposed layer | Derivation and held-out evidence | Single-cell context | Analytical role |
| --- | --- | --- | --- | --- |
| STING-core | Entry anchor | Derivation lesion-enriched; GSE121212 paired delta 1.288 (95% CI 1.063-1.513) | Positive lesional shifts in KC and myeloid/DC | Reference axis; not biochemical activation |
| Proximal bridge | Proximal core | Strong derivation enrichment/coupling; held-out coupling rho 0.767 | Positive across several broad compartments | Target-proximal research layer |
| IFN-responsive | Primary branch | Strong derivation evidence; held-out paired delta 1.478 and coupling rho 0.737 | Broad positive lesional distribution; day-14 contraction | Response-facing core branch |
| NF-κB broad innate | Secondary innate | Positive derivation effect; held-out STING-high effect positive but weaker | Compartment-dependent | Secondary inflammatory branch |
| Keratinocyte effector | Effector | Treatment-reversible; held-out lesion enrichment and coupling rho 0.565 | Strong lesional KC and myeloid/DC shifts | Amplification/pharmacodynamic output |
| Myeloid/neutrophil | Effector | Positive lesion and treatment signals; held-out STING-high effect positive | S100-linked activity across compartments | Effector/context-sensitive output |
| Immune proxy | Context | Weak derivation coupling but positive held-out context activity | Broad immune-representation context | Context modifier; not target branch |
| Keratinocyte differentiation | Context | Weak derivation signal; held-out STING-high effect strongly negative | Lower in lesional KC | Differentiation context control |

Note: Evidence integrates the derivation cohorts, held-out GSE121212 replication and GSE228421 broad-cell-type context analysis. The held-out verdict was partial replication: the lesion anchor and proximal/IFN-dominant coupling structure transported, whereas complete STING-high branch ordering did not. Full statistics are provided in Online Resources 3, 5 and 6.

Keratinocyte-effector, NF-κB broad innate and myeloid/neutrophil proxy branches formed secondary or downstream layers. The strongest pairwise internal edge linked IFN-responsive and proximal-bridge branches. This organization supports a layered branch architecture rather than a flat inflammatory score.

### Held-out GSE121212 replicates the lesion anchor and proximal/IFN-dominant coupling structure

In 27 matched psoriasis pairs, the primary TMM-log2CPM lane showed lesional-minus-nonlesional effects of 1.288 for STING-core (95% CI 1.063 to 1.513; P = 6.56 x 10^-12), 1.478 for IFN-responsive activity (95% CI 1.103 to 1.852; P = 1.34 x 10^-8) and 1.188 for proximal-only STING (95% CI 1.031 to 1.345; P = 1.12 x 10^-14) (Fig. 3A; Online Resource 6). The prespecified Anchor component therefore passed.

**Figure 3.**
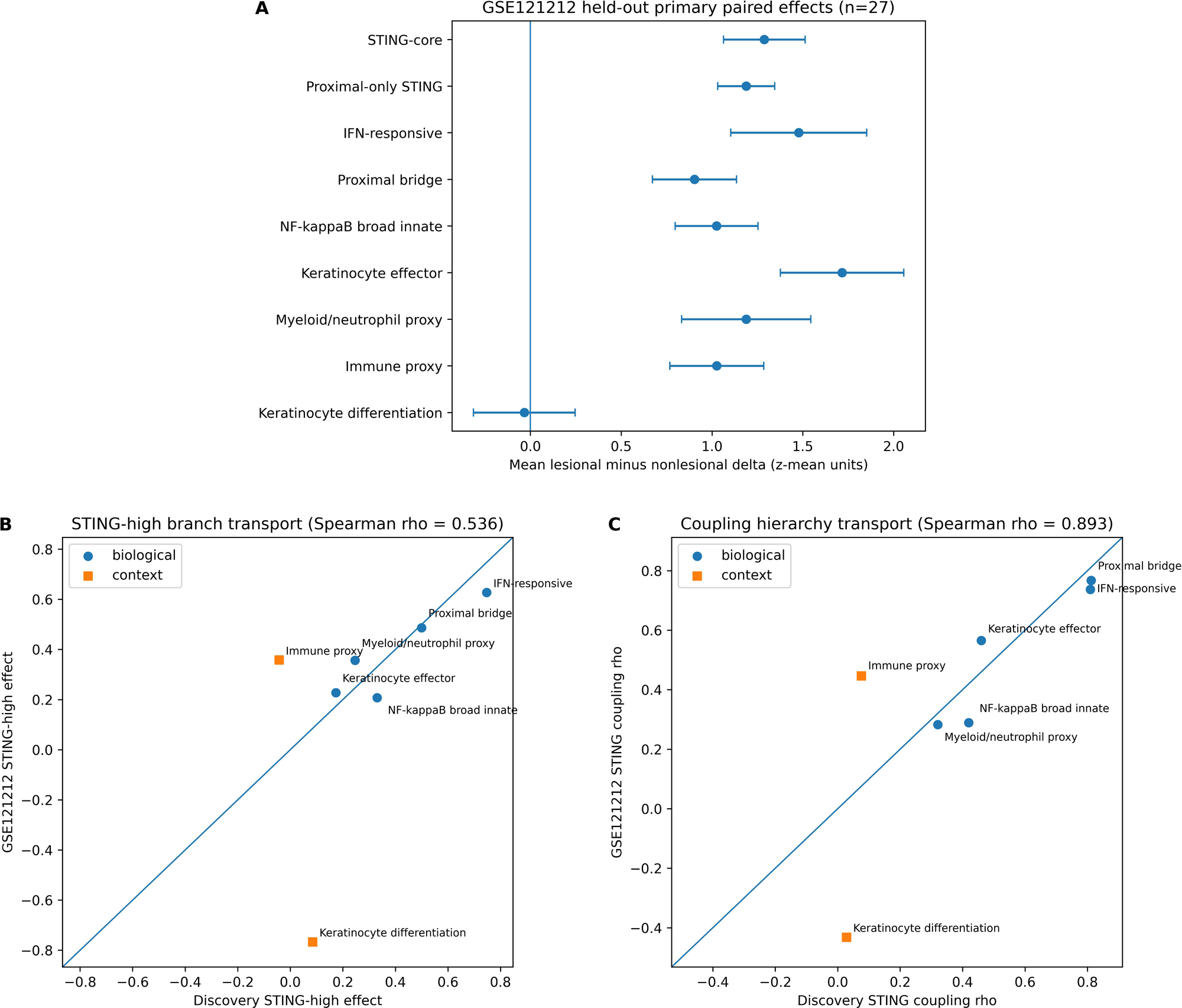
Held-out GSE121212 replication. (A) Primary TMM-log2CPM paired lesional-minus-nonlesional effects in 27 matched psoriasis pairs. Points show mean patient-level deltas and bars show 95% confidence intervals. (B) Discovery versus held-out STING-high effects across the seven frozen downstream branches; rank concordance was moderate (Spearman rho = 0.536). (C) Discovery versus held-out lesional STING-core coupling across the same branches; rank concordance was strong (rho = 0.893, P = 0.0068). The prespecified verdict was Anchor PASS, Architecture PARTIAL and Overlap-robust PASS.

Among 28 lesional samples, proximal bridge and IFN-responsive branches retained the strongest STING-core coupling (rho = 0.767 and 0.737, respectively). The seven-branch coupling vector was strongly concordant with the frozen discovery vector (rho = 0.893, P = 0.0068; Fig. 3C). STING-high effects were positive for all five biological branches, but the complete effect ordering was only moderately concordant with discovery (rho = 0.536; Fig. 3B), and the biological-versus-context criterion was not met because immune-proxy activity was comparatively high and keratinocyte differentiation was strongly negative.

The frozen component verdicts were Anchor PASS, Architecture PARTIAL and Overlap-robust PASS, yielding PARTIAL_REPLICATION overall. The legacy raw-count lane retained the anchor but failed architecture transport and broadly increased context-branch coupling. Accordingly, the TMM-log2CPM result is authoritative and detailed branch ordering is interpreted as preprocessing-sensitive (Online Resource 6).

### Treatment-facing bulk cohorts show directional branch reversibility

In GSE117239, PASI75 responders had greater week-12 reductions in interferon and STING-core modules than nonresponders (Fig. 4; Online Resource 3). Responder lesions showed significant decline in IFN-responsive, proximal-bridge, NF-κB broad innate, myeloid/neutrophil and keratinocyte-effector branches. The largest declines were observed for IFN-responsive and keratinocyte-effector branches.

**Figure 4.**
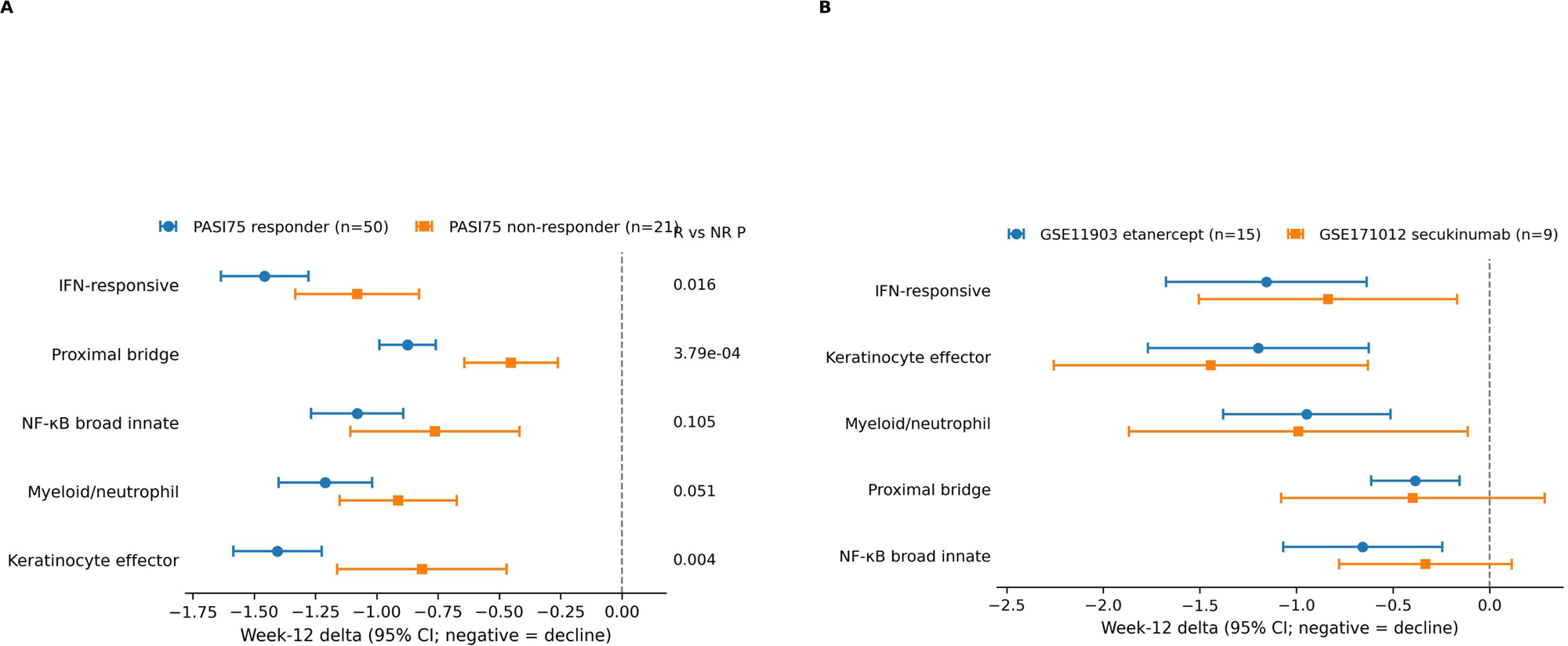
Treatment-linked branch remodeling in bulk cohorts. Treatment-facing bulk datasets show directional contraction of IFN-responsive, proximal-bridge and effector-associated branches, especially in responder lesions. These analyses support pharmacodynamic remodeling of the lesion-state architecture rather than direct causal targeting of STING.

External week-12 analyses in GSE11903 and GSE171012 showed additional directional contraction of major downstream branches. In GSE11903, STING-core, proximal bridge, IFN-responsive, NF-κB broad innate, keratinocyte effector and myeloid/neutrophil branches all declined at week 12. In GSE171012, IFN-responsive, keratinocyte effector and myeloid/neutrophil branches declined significantly, while proximal bridge was directionally negative but weaker. These findings support pharmacodynamic branch remodeling but do not establish treatment equivalence or causal targeting of STING.

### Robustness analyses support structure beyond gene overlap and tissue composition

The proximal-only STING reference attenuated but did not remove the paired lesion enrichment or interferon coupling, indicating that the STING-IFN relationship was not created solely by shared downstream interferon-output genes (Fig. 5; Online Resource 3). Composition-aware models retained major STING-associated branch relationships after adjustment for immune-proxy and keratinocyte-context scores.

**Figure 5.**
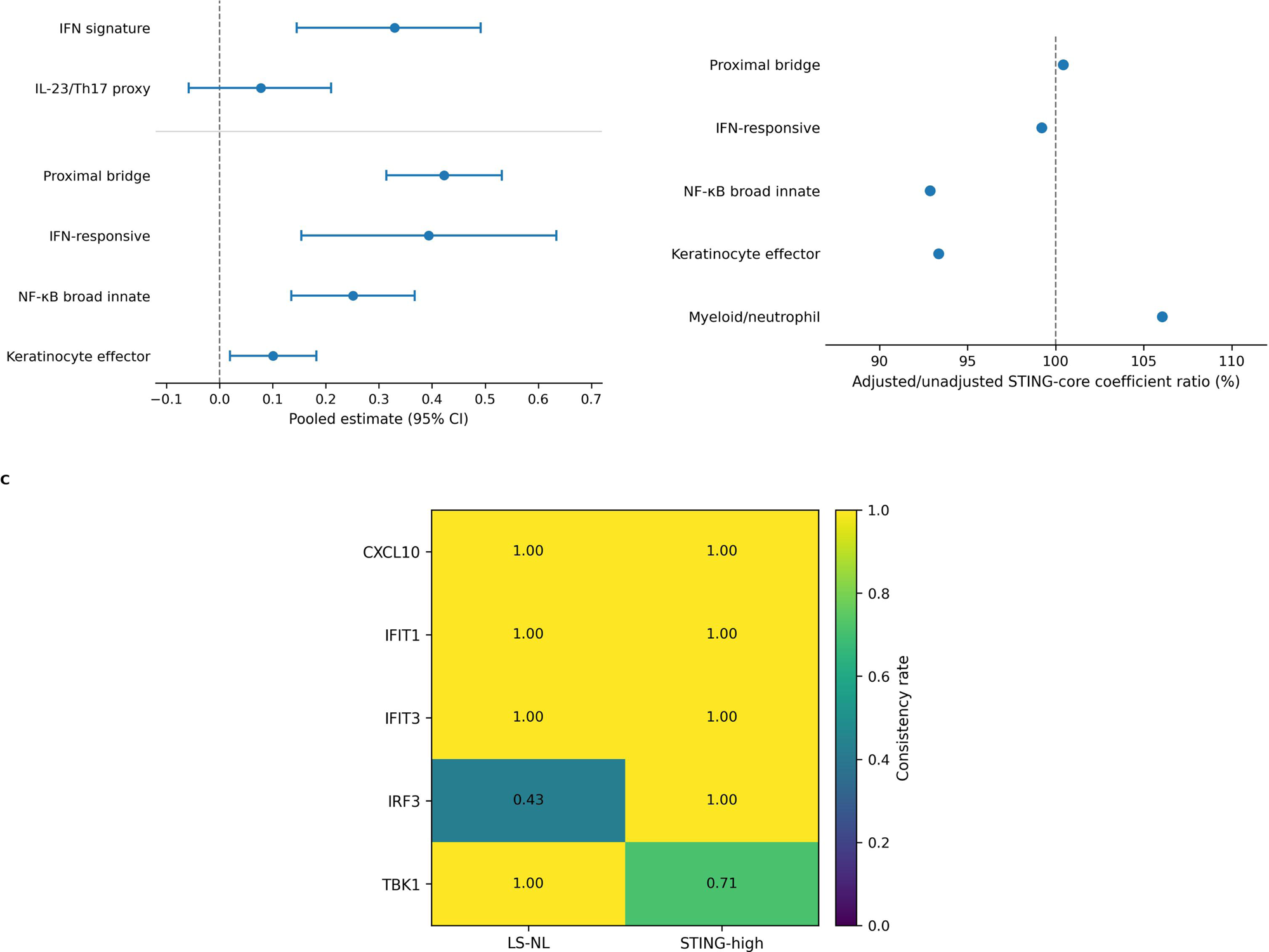
Robustness and gene-level anchoring. Proximal-only STING sensitivity, composition-aware adjustment and representative-gene analyses support the branch structure beyond downstream gene overlap and trivial tissue-composition effects.

Representative-gene analyses further anchored the framework: CXCL10, IFIT1, IFIT3 and TBK1 were positive in all seven paired lesion comparisons, whereas IRF3 was inconsistent in bulk lesion comparisons but positive in STING-high analyses, compatible with a regulatory component whose transcript-level signal is context dependent.

### Single-cell analysis shows broad-cell-type-stratified distribution and major-composition robustness

The GSE228421 single-cell extension retained 166,954 cells from 17 samples after independent per-sample raw-droplet filtering. Manual annotation identified 158,503 primary cells after exclusion of 8,451 low-confidence mixed cells.

Patient-level paired and longitudinal inference was restricted to four complete patients. The major included compartments were keratinocyte (68,186 cells), fibroblast (29,440), myeloid/DC (18,867), endothelial (18,478), pericyte/smooth-muscle

(12,923), T/NK (4,266), melanocyte (3,488), mast cell (2,166) and B/plasma cell (689) (Fig. 6; Online Resource 4). Baseline paired broad-cell-type analysis used P1-P4 only. In keratinocytes, STING-core, proximal bridge, IFN-responsive, keratinocyte-effector and myeloid/neutrophil-proxy scores were higher in lesional than matched nonlesional skin in all four patients, whereas keratinocyte differentiation was lower. Myeloid/DC cells showed concordant positive lesional shifts in the same major inflammatory branches. Fibroblast and pericyte/smooth-muscle compartments also showed positive IFN/STING-linked shifts. Persistence within several broad annotated compartments argues against an explanation based solely on shifts among major cell-type proportions, while leaving finer cellular substates and within-compartment composition unresolved (Online Resource 5).

**Figure 6.**
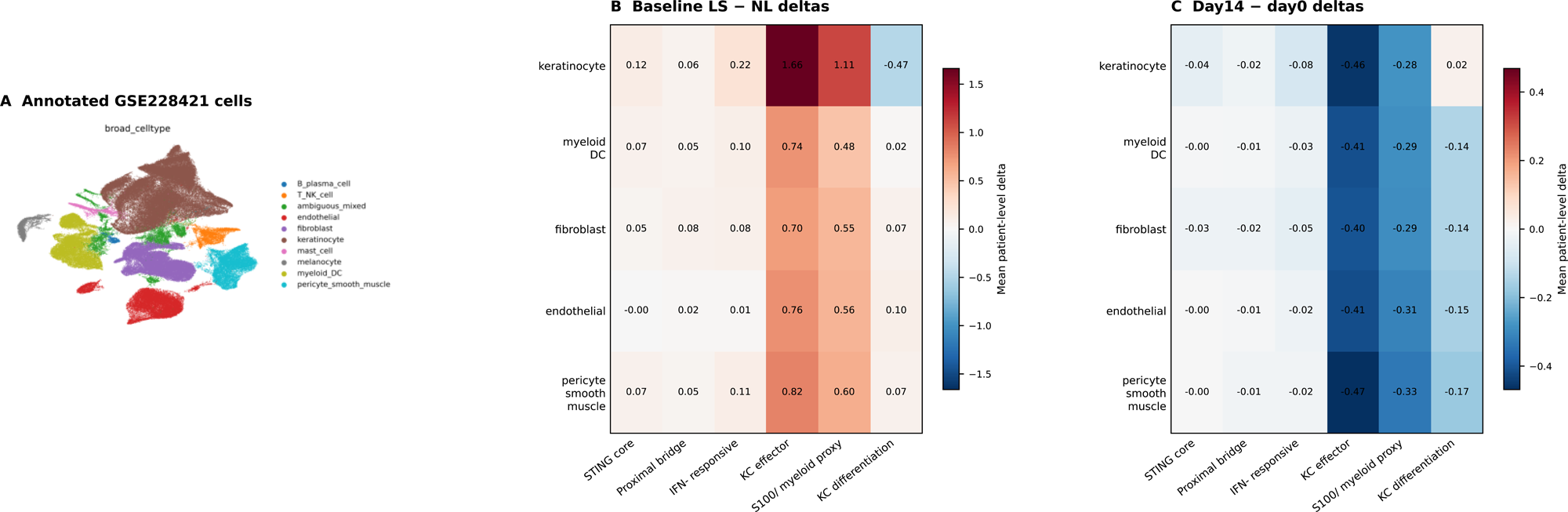
Single-cell context and major-composition robustness. Annotated GSE228421 cells show the distribution of frozen branches across broad cellular compartments and patient-level lesional/nonlesional and day-14 trajectories. Patient-level paired and longitudinal inference was restricted to four complete patients. Persistence within several broad cell types argues against an explanation based solely on major cell-type proportion shifts, but does not identify a unique cell of origin or exclude finer within-compartment composition.

During risankizumab treatment, day-14 lesional trajectories showed directional contraction of IFN/STING-linked and selected effector-associated activity in several broad compartments. Keratinocyte IFN-responsive and STING-core scores decreased in all four longitudinal patients, and myeloid/DC and fibroblast summaries showed similar directional contraction. These descriptive trajectories were not uniform across all branches or compartments and therefore support within-compartment remodeling rather than complete reversal of the bulk hierarchy (Online Resource 5).

## Discussion

This study defines and externally tests a branch-resolved IFN/STING-linked lesion-state architecture in human psoriasis. The main contribution is not the discovery of a previously unknown psoriasis pathway. Interferon, NF-κB, keratinocyte effector, myeloid/neutrophil and differentiation biology are established components of psoriasis [1–23]. The added value is that these components were organized into a frozen architecture with distinguishable proximal, response-facing, effector and context branches, followed by an untouched bulk replication that was allowed to fail under prespecified criteria. This converts established inflammatory programmes from a parallel list of enrichments into a falsifiable architecture whose generalization boundary can be measured.

The held-out result clarifies which properties are transportable. The IFN/STING lesion anchor, proximal-only sensitivity and proximal/IFN-dominant coupling structure replicated strongly. In contrast, the complete STING-high magnitude ordering was only partially transportable, and context branches were not invariant. This boundary is scientifically informative: the architecture has a stable core, but its broader tissue-expression ordering can vary with assay platform, patient recruitment, anatomical sampling, clinical context and RNA-sequencing preprocessing. The legacy raw-count sensitivity cannot override the primary lane because it inflated coupling across both biological and context branches.

The GSE78097 severity-context result reinforces this distinction between lesion presence and severity. Severe lesions showed lower IFN-responsive and myeloid/neutrophil scores, a lower STING-core point estimate, and higher keratinocyte-proxy activity than mild lesions. This non-monotonic pattern may reflect differences in tissue composition, lesion stage, anatomical sampling or molecular-state mixture, but the available metadata do not distinguish among these possibilities. The IFN/STING axis should therefore be interpreted as a reproducible lesion-state anchor rather than a linear surrogate for clinical severity.

With only four complete inferential patients, the single-cell analysis addresses a narrower question. Frozen branch activity persisted within keratinocyte, myeloid/DC and stromal compartments, arguing against an explanation based solely on shifts among major annotated cell types. It does not identify a unique cell of origin, independently replicate the complete bulk hierarchy or exclude finer substate composition. The phrase cellular-context analysis therefore better reflects its evidentiary role than independent single-cell validation.

The treatment-facing analyses should be interpreted as pharmacodynamic remodeling, not as causal proof. Ustekinumab, etanercept, secukinumab and risankizumab act at different points in the inflammatory network. The shared observation that IFN/STING-linked and effector-associated branches contract during treatment suggests that the architecture tracks lesion resolution. In the risankizumab single-cell cohort, early contraction of IFN/STING-linked branches after IL-23 blockade is best interpreted as indirect lesion-state remodeling rather than direct STING inhibition, consistent with independent single-cell evidence that IL-23 inhibition reshapes lesional cell states without a single dominant effector population [45]. One plausible explanation is that IL-23/Th17-axis blockade reduces keratinocyte and myeloid inflammatory loops that help sustain interferon-associated branch activity, but the present dataset does not establish the molecular sequence of this cross-axis effect. It does not show that STING is the direct therapeutic target, nor that IFN/STING-linked activity is upstream of all other branches.

The framework has a bounded translational pathway. A compact RNA panel could separate target-proximal or bridge measurements from downstream response-facing outputs, in the same spirit as recent transcriptomic-score approaches to psoriasis severity and treatment response [9]. Such a panel may be useful in biopsy-based trials to monitor whether a plaque is moving away from an inflammatory lesion state. However, clinical use would require prospective thresholding, repeatability assessment, pre-specified endpoint testing and comparison with existing clinical and molecular markers. The present study defines the architecture and its validation route; it does not provide a diagnostic or treatment-selection test. The incomplete transportability of the broader branch ordering may reflect more than technical cross-cohort variation.

One possibility is that cohorts contain different mixtures of latent molecular lesion states that share a stable IFN/STING and proximal core but differ in the deployment of innate, keratinocyte, myeloid and context programmes. The present study cannot distinguish such biological heterogeneity from platform, sampling or other dataset-domain effects.

Nevertheless, this combination of a reproducible core and variable peripheral configuration provides a restrained rationale for future multidimensional lesion-state modeling, including virtual-lesion approaches in which branch weights and treatment trajectories are learned rather than assumed to follow a universal fixed hierarchy.

Several limitations define the boundary of inference. First, the study is observational and based on public cohorts with platform, sampling and treatment differences. Second, transcriptomic activity does not establish biochemical STING activation; no STING protein, phosphorylation, localization, cGAS activity, phospho-TBK1 or phospho-IRF3 measurements were available. Third, GSE121212 is a retrospective held-out analysis rather than a prospectively collected cohort, and the architecture verdict was partial rather than complete. Fourth, detailed branch ordering was sensitive to RNA-sequencing preprocessing. Fifth, the single-cell analysis used four complete inferential patients and broad annotations; cells were not treated as independent replicates, and finer within-compartment states remain unresolved. Sixth, overlapping genes such as S100A8 and S100A9 require branch-level rather than cell-identity interpretation.

Finally, STING-high and any proposed panel thresholds require prospective validation.

Within these limits, a branch-resolved architecture provides more information than a single inflammatory score. The held-out analysis supports a stable IFN/STING and proximal core while explicitly rejecting the assumption that every secondary or context branch has invariant ordering. This bounded framework is suitable for hypothesis-driven pharmacodynamic monitoring and future model development, but not yet for direct clinical decision-making.

## Conclusions

Human psoriasis plaques exhibit a reproducible IFN/STING-linked lesion anchor embedded within a broader branch-resolved inflammatory architecture. An untouched bulk RNA-sequencing cohort replicated the lesion anchor and proximal/IFN-dominant coupling structure, while the complete seven-branch STING-high ordering showed only partial transportability. Single-cell analysis supports broad-cell-type-stratified distribution and robustness against a major-cell-proportion-only explanation. Prospective cohorts and orthogonal protein, spatial or functional measurements remain necessary before mechanistic or clinical claims can be made.

## Declarations

### Ethics approval

This secondary analysis used only public, de-identified datasets and required no institutional review board approval under local regulations.

### Consent to participate

Not applicable.

### Consent for publication

Not applicable.

### Data availability

All analysed transcriptomic datasets are publicly available through GEO under the accession numbers cited in the reference list. Summary tables, frozen-design documents, figure-source matrices and workflow documentation are included in the Online Resources and companion machine-readable packages or are available from the corresponding author during review.

### Code availability

Curated analysis scripts and execution notes are available from the corresponding author during peer review and will be deposited in a suitable public repository if required before publication.

### Author contributions

Conceptualization, T.-L.C. and H.-W.S.; methodology, T.-L.C. and H.-Y.S.; software and formal analysis, T.-L.C. and H.-Y.S.; investigation, T.-L.C. and H.-Y.S.; resources, H.-W.S. and Z.-H.W.; writing - original draft, T.-L.C.; writing - review and editing, all authors; supervision, H.-W.S. and Z.-H.W.; project administration, H.- W.S. All authors reviewed and approved the final manuscript.

### Artificial intelligence-assisted writing disclosure

Artificial intelligence was used only as an editorial tool to improve English language and readability. No AI system was used for study design, data selection, analysis, interpretation, figure or table generation, or conclusions. All edited text was reviewed and approved by the authors.

## Supporting information

supplemental informations

## Acknowledgements

The authors thank the investigators who generated and deposited the public datasets reanalysed in this study.

## Online Resource captions

**Online Resource 1.** Cohort inventory and analysis roles. Derivation bulk cohorts, held-out GSE121212, treatment-facing cohorts and GSE228421 cellular-context design.

**Online Resource 2.** Frozen module and branch definitions. Primary modules, downstream branches, gene sets and single-cell gene availability.

**Online Resource 3.** Bulk transcriptomic branch architecture. Paired lesion effects, STING coupling, STING-high differences, treatment reversibility and robustness statistics.

**Online Resource 4.** GSE228421 single-cell quality control and annotation. Raw droplet filtering, per-sample cell counts, manual annotation and mixed-cluster exclusion.

**Online Resource 5.** Broad-cell-type-stratified branch context analysis. Baseline lesional-minus-nonlesional and day14-minus-day0 matrices and conservative interpretation guide.

**Online Resource 6.** GSE121212 held-out bulk replication. Frozen design, gene coverage, primary paired effects, STING-high and coupling transport, verdicts and preprocessing sensitivity.

**Online Resource 7.** Reproducibility and reporting notes. Script manifest, data and code availability, checksums, IL-23/Th17 source-authority reconciliation and artificial-intelligence disclosure.

