## supplemental informations for "Branch-resolved IFN/STING-linked lesion-state architecture in psoriasis: multi-cohort derivation, held-out bulk replication and single-cell context analysis": README_IR_v4_1_1_clean_machine_readable_tables.md

# IR v4.1.1 clean machine-readable table package

This package accompanies **Branch-resolved IFN/STING-linked lesion-state architecture in psoriasis: multi-cohort derivation, held-out bulk replication and single-cell context analysis**.

## Additions in v4.1.1
- Held-out GSE121212 frozen-design and primary TMM-log2CPM replication outputs.
- Publication-facing Online Resource Tables S26-S31.
- Primary-versus-legacy preprocessing sensitivity sources.
- IL-23/Th17 source-authority reconciliation.

The publication-authority IL-23/Th17 proxy contains 11 genes. A same-name 8-gene panel from a separate virtual-lesion/module-score workflow is preserved only in raw provenance and is excluded from publication-facing GSE121212 results.
