## supplemental informations for "Branch-resolved IFN/STING-linked lesion-state architecture in psoriasis: multi-cohort derivation, held-out bulk replication and single-cell context analysis": V4_1_1_EDITORIAL_UPDATE_NOTE.md

# v4.1.1 editorial update note

The numerical content of this machine-readable package is unchanged from the frozen IR v4.0/V6F authority. Version 4.1.1 changes manuscript wording, restores the GSE78097 main-text interpretation, renumbers main figures by first citation, and corrects Figure S6 connector colours. Selected internal filenames retain `v4_0` identifiers to preserve source provenance and prevent the editorial update from being mistaken for a numerical re-analysis.
