## Supplementary figures and images for "Branch-resolved IFN/STING-linked lesion-state architecture in psoriasis: multi-cohort derivation, held-out bulk replication and single-cell context analysis"

### Figure s1

**Online Resource Figure S1. Expanded representative-gene anchoring**

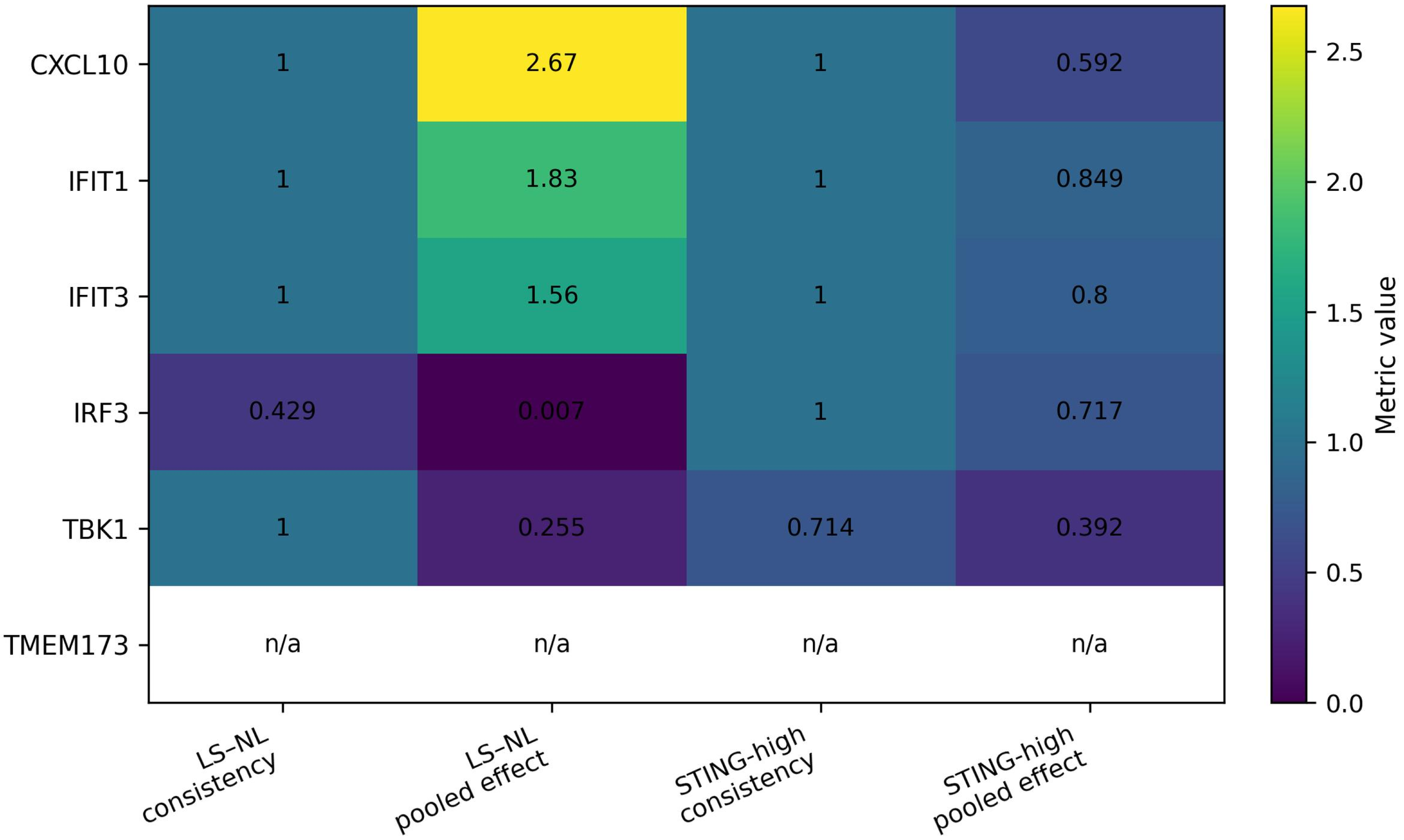

### Figure s2

**Online Resource Figure S2. Severity and composition context**

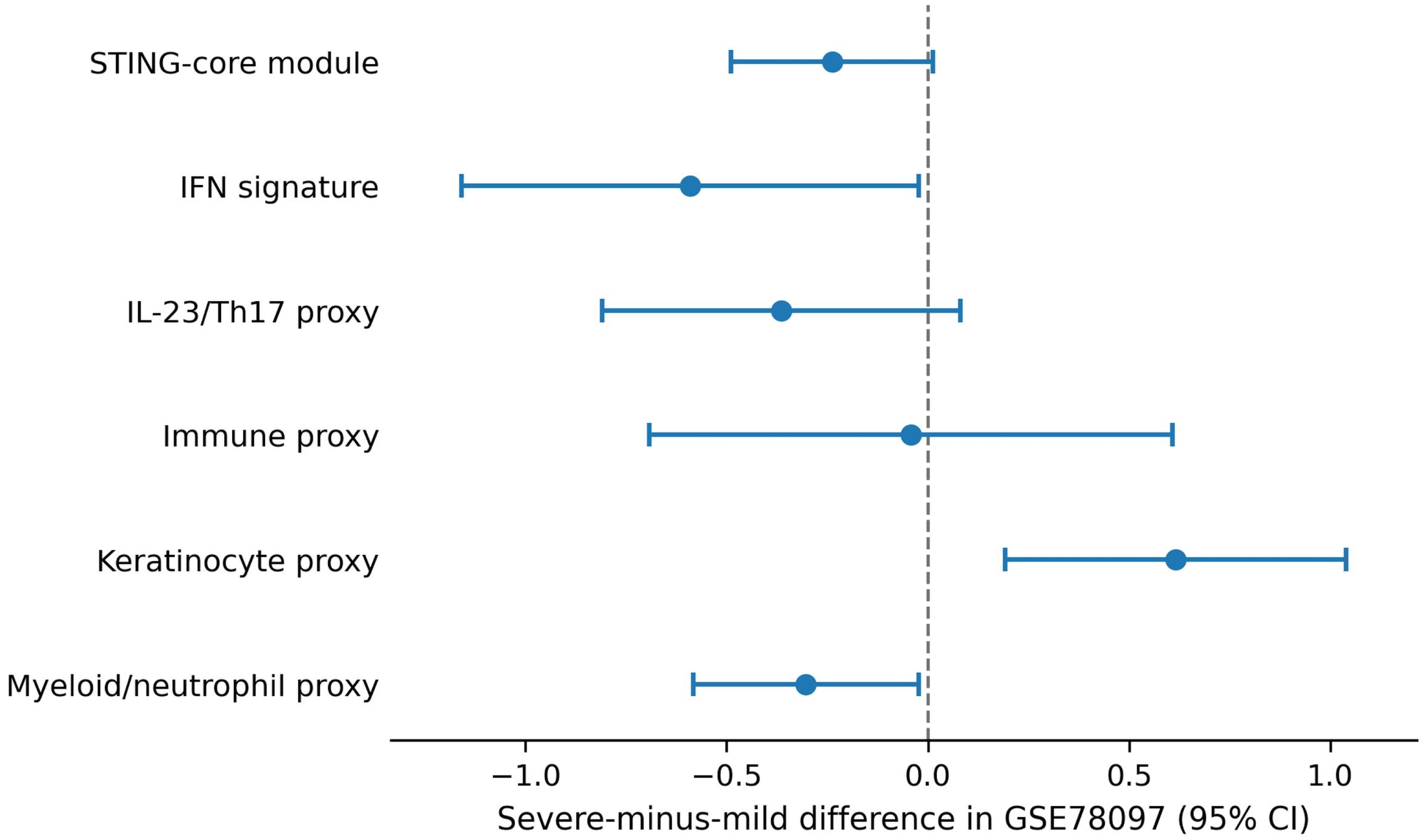

### Figure s4

#### Online Resource Figure S4. Full baseline LS − NL celltype × branch matrix

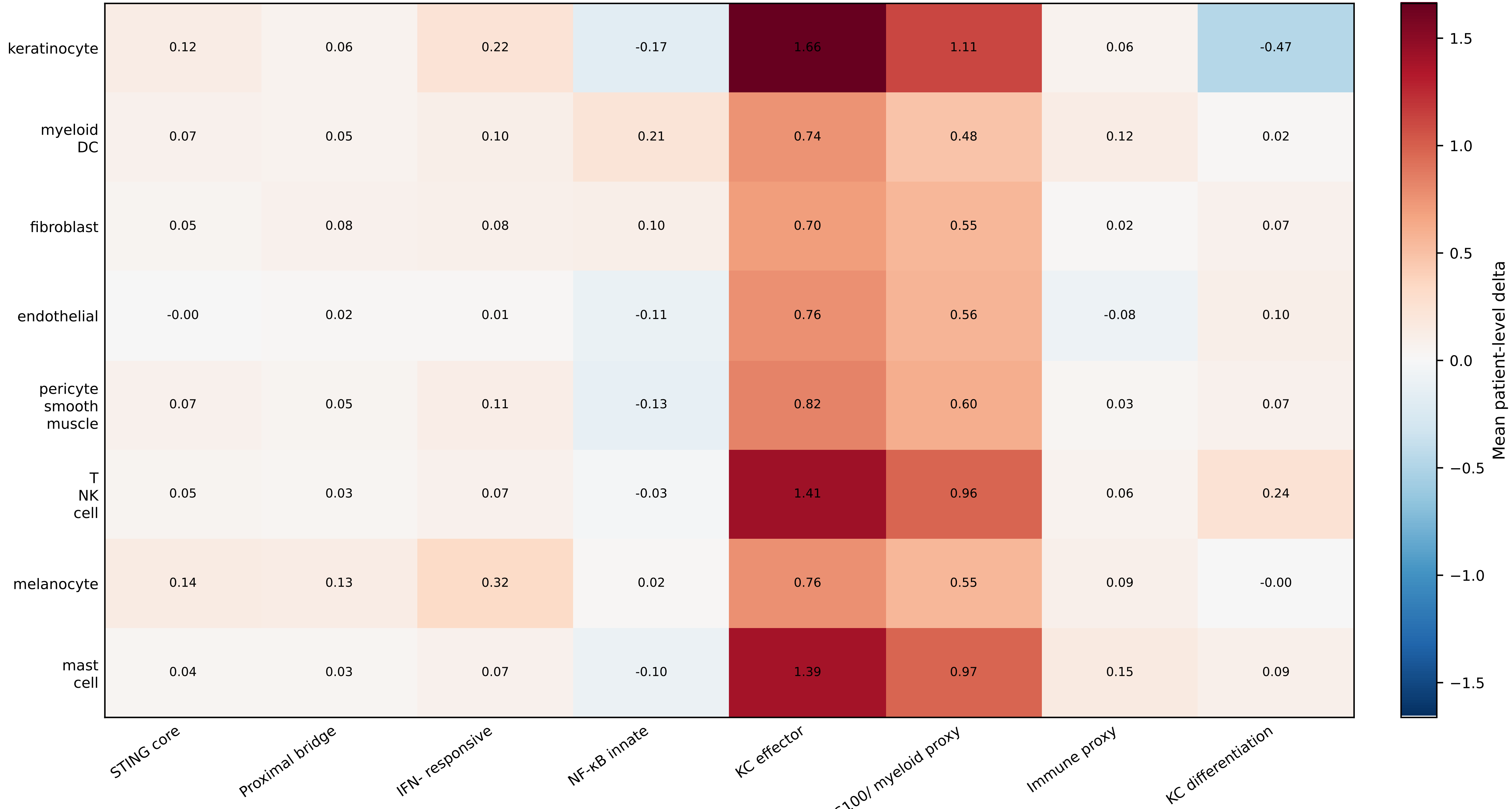

### Figure s5

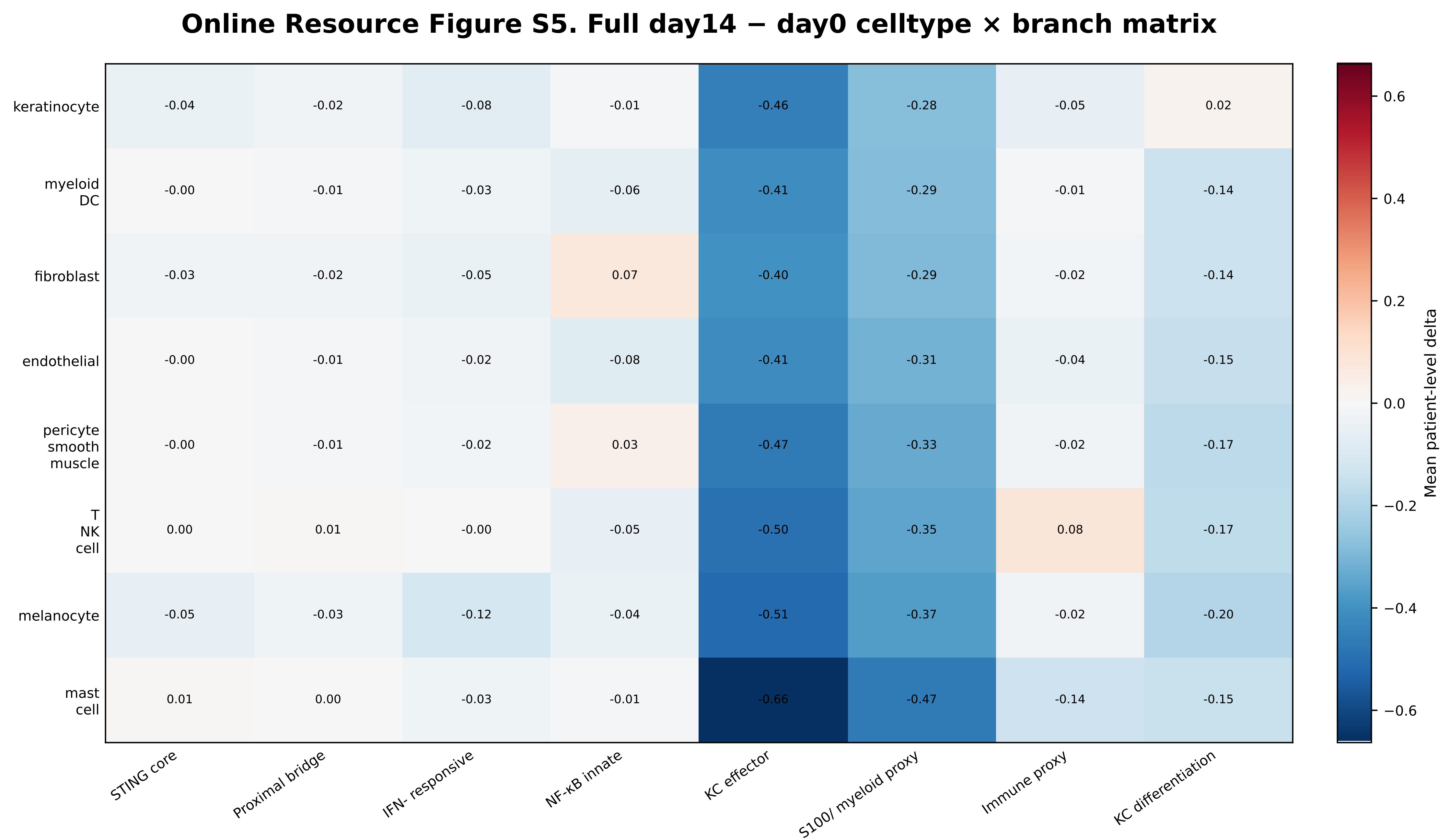

### Figure s6

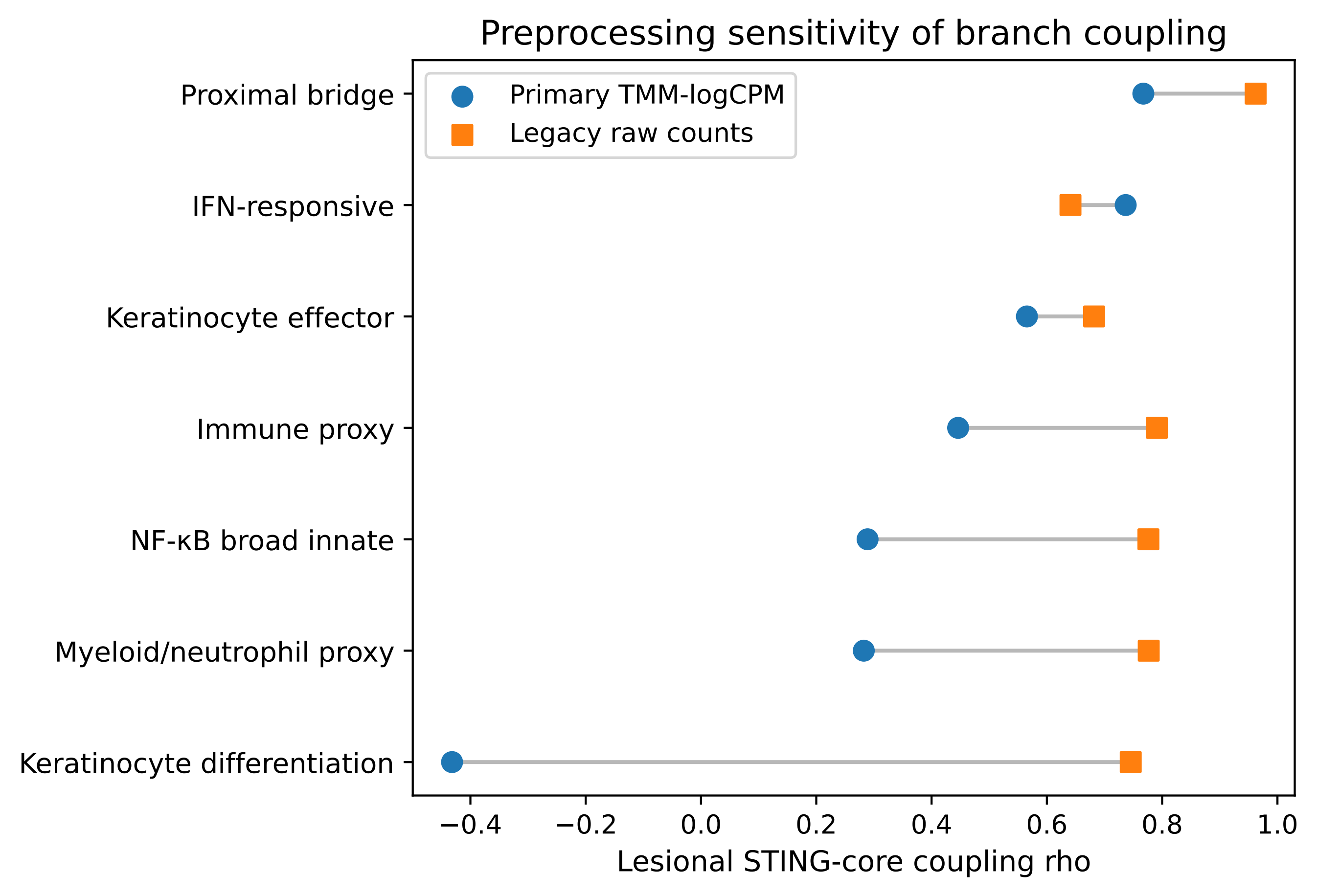
