## Supplementary material for "Branch-resolved IFN/STING-linked lesion-state architecture in psoriasis: multi-cohort derivation, held-out bulk replication and single-cell context analysis": Figure s3

### Online Resource Figure S3. GSE228421 annotation and primary-inference QC

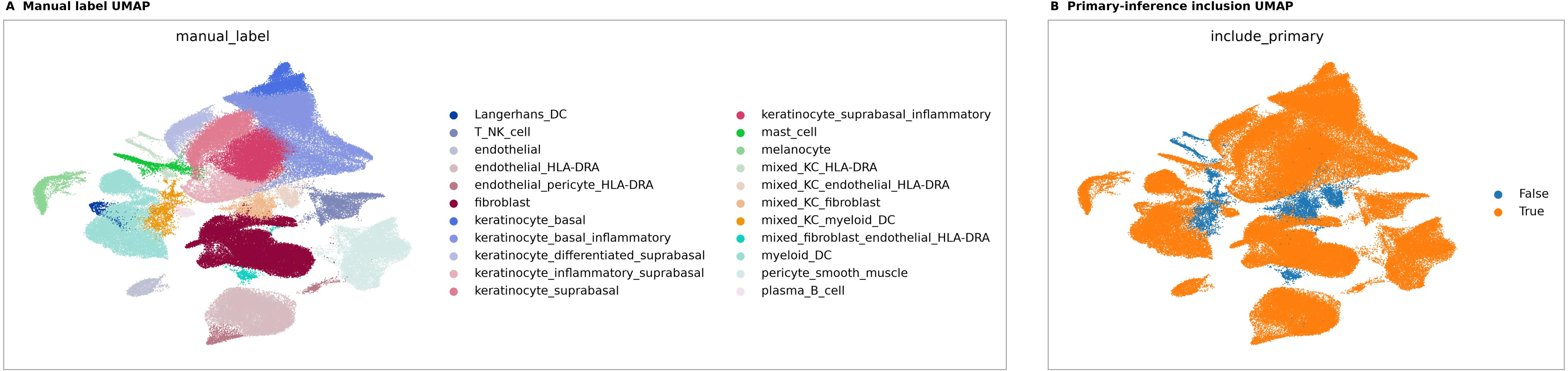


Panels preserve the original source UMAP images and color legends without cropping. Manual labels show broad annotation groups; primary-inference inclusion separates retained cells from excluded ambiguous/mixed cells.
